# Mechanism and consequences of herpes simplex virus 1-mediated regulation of host mRNA alternative polyadenylation

**DOI:** 10.1101/2020.11.26.399626

**Authors:** Xiuye Wang, Liang Liu, Adam W. Whisnant, Thomas Hennig, Lara Djakovic, Nabila Haque, Cindy Bach, Rozanne M. Sandri-Goldin, Florian Erhard, Caroline C. Friedel, Lars Dölken, Yongsheng Shi

## Abstract

Eukaryotic gene expression is extensively regulated by cellular stress and pathogen infections. We have previously shown that herpes simplex virus 1 (HSV-1) and several cellular stresses cause widespread disruption of transcription termination (DoTT) of RNA polymerase II (RNAPII) in host genes and that the viral immediate early factor ICP27 plays an important role in HSV-1-induced DoTT. Here, we show that HSV-1 infection also leads to widespread changes in alternative polyadenylation (APA) of host mRNAs. In the majority of cases, polyadenylation shifts to upstream poly(A) sites (PAS), including many intronic PAS. Mechanistically, ICP27 contributes to HSV-1-mediated APA regulation. HSV-1- and ICP27-induced activation of intronic PAS is sequence-dependent and does not involve general inhibition of U1 snRNP. HSV1-induced intronic polyadenylation is accompanied by early termination of RNAPII. Finally, HSV-1-induced mRNAs polyadenylated at intronic PAS are exported into the cytoplasm while APA isoforms with extended 3’ UTRs are sequestered in the nuclei, both preventing the expression of the full-length gene products. Together with other recent studies, our results suggest that viral infection and cellular stresses induce a multi-faceted host shutoff response that includes DoTT and changes in APA profiles.

## Introduction

The 3’ ends of the vast majority of eukaryotic mRNAs are formed through cleavage and polyadenylation [1–3]. In mammals, poly(A) sites (PAS) are defined by several cis-elements, including the AAUAAA hexamer, the U/GU-rich downstream element, and other auxiliary sequences. These sequences recruit RNA 3’ processing factors CPSF, CstF, CFIm, CFIIm, and the poly(A) polymerase to form the 3’ processing complex. RNA 3’ processing occurs co-transcriptionally and it plays an essential role not only in RNA biogenesis, but also in transcription termination by RNA polymerase II (RNAPII) [4–6]. According to the “allosteric model” of transcription termination, the transcription machinery undergoes a transformation upon passing through a PAS, which primes RNAPII for termination. Alternatively, the “torpedo model” posits that the unprotected 5’ end of RNA generated by the 3’ processing cleavage step is recognized by the exoribonuclease Xrn2. Xrn2-mediated degradation of the nascent RNA ultimately leads to transcription termination. Thus, in both models, RNA 3’ processing plays a central role in transcription termination.

RNA 3’ processing also plays an important role in gene regulation. The transcripts of over 70% of human genes can be cleaved and polyadenylated at multiple alternative PAS, a process called alternative polyadenylation (APA) [7–9]. Different APA isoforms from the same gene may encode distinct proteins and/or contain different 3’ untranslated regions (UTRs). 3’ UTRs are hot spots for regulation: they harbor target sites for microRNAs, binding sites for RNA-binding proteins, RNA localization signals, and they can function as protein assembly platforms. Thus, APA isoforms from the same gene could be differentially regulated. Recent studies have provided evidence that APA plays important roles in a wide variety of biological processes and aberrant APA regulation has been linked to a number of diseases, including cancer and neurological disorders [10]. Many APA regulators have been identified, including the core RNA 3’ processing factors, splicing factors, and RNA-binding proteins [11]. For example, U1 snRNP has been shown to inhibit premature cleavage/polyadenylation at intronic PAS, thereby protecting transcript integrity globally [12]. Despite recent progress, however, the regulatory mechanisms and functional consequences of APA remain poorly understood.

Both RNA 3’ processing and transcription termination are highly regulated. For example, we have previously shown that HSV-1 infection leads to a widespread disruption of transcription termination (DoTT) [13][14]. Influenza virus (IAV) was reported to elicit a similar response [15]. The Steitz lab observed a transcription termination defect in cells exposed to salt/osmotic stress that leads to the production of transcripts downstream of genes (DoGs) [16]. A comparative analysis showed that virus-induced DoTT and stress-induced DoGs are highly related [17]. Although the mechanism for DoTT/DoGs remains unclear, we have recently shown that the viral immediate early factor ICP27 contributes to HSV-1-induced DoTT by directly binding to the RNA 3’ processing factor CPSF and inhibiting the cleavage step [13]. Meanwhile, several groups reported that virus infections, such as the human cytomegalovirus (HCMV) and vesicular stomatitis virus (VSV), or stress can induce global APA changes [18–20]. However, the relationship between virus- or stress-induced APA and DoTT/DoG remains unclear. In this study, we integrated time-resolved global APA profiling, nascent RNA sequencing, cell fractionation and RNA sequencing data in HSV-1-infected cells to elucidate the scope, mechanism, and functional impact of virus-induced APA changes and DoTT.

## Results

### HSV-1 infection induces widespread and dynamic APA changes

To determine if and how the global APA profile of host genes is altered during HSV-1 infection, we performed PAS-seq analysis of HeLa cells at 0, 2, 6, and 12 hours post-infection (hpi). PAS-seq is a method developed in our laboratory for quantitatively mapping RNA poly(A) junctions and has been used extensively for profiling global APA [21,22]. By comparing the APA profiles of cells at 0 and 12 hpi, we detected significant APA changes in 1,050 genes (FDR < 0.05, impacting at least 15% of transcripts, see Methods for details). In 745 genes (71%), polyadenylation shifted to proximal PAS (Distal-to-Proximal or DtoP) and 305 (29%) showed changes in the opposite direction (Proximal-to-Distal or PtoD) (Fig. 1A). A significant portion of these changes involved intronic PAS. For example, among the DtoP changes, 331 (44%) shifted to a proximal intronic PAS (DtoP_Intron) while 32% of PtoD genes shifted from a proximal intronic PAS to a PAS in the 3’ UTR (PtoD_Intron, Fig. 1A). APA profile changes could be due to differential PAS selection during transcription, differential degradation of APA isoforms, a selective loss of proximal or distal PAS due to read-through transcription, or a combination of all factors. To begin to understand the cause of the observed APA changes in HSV-1-infected cells, we compared the PAS-seq reads (normalized by sequencing depths of the host transcriptomes) at proximal and distal PAS of genes that displayed significant APA changes. As shown in Fig. 1B, DtoP changes are accompanied by a relative increase in proximal PAS reads and a relative decrease in distal PAS reads. Conversely, PtoD changes involving only UTR PAS are accompanied by the opposite changes. In addition to ratios, DtoP changes are accompanied by a net increase in the read counts at proximal PAS and a net decrease at the distal PAS, while the opposite changes were observed for PtoD shifts (Supplemental Fig. 1). These results provided evidence that the HSV-1-induced APA changes are not caused solely by preferential degradation or loss of specific APA isoforms, but may require a shift in PAS usage. The underlying mechanism will be further addressed below.

**Figure 1.**
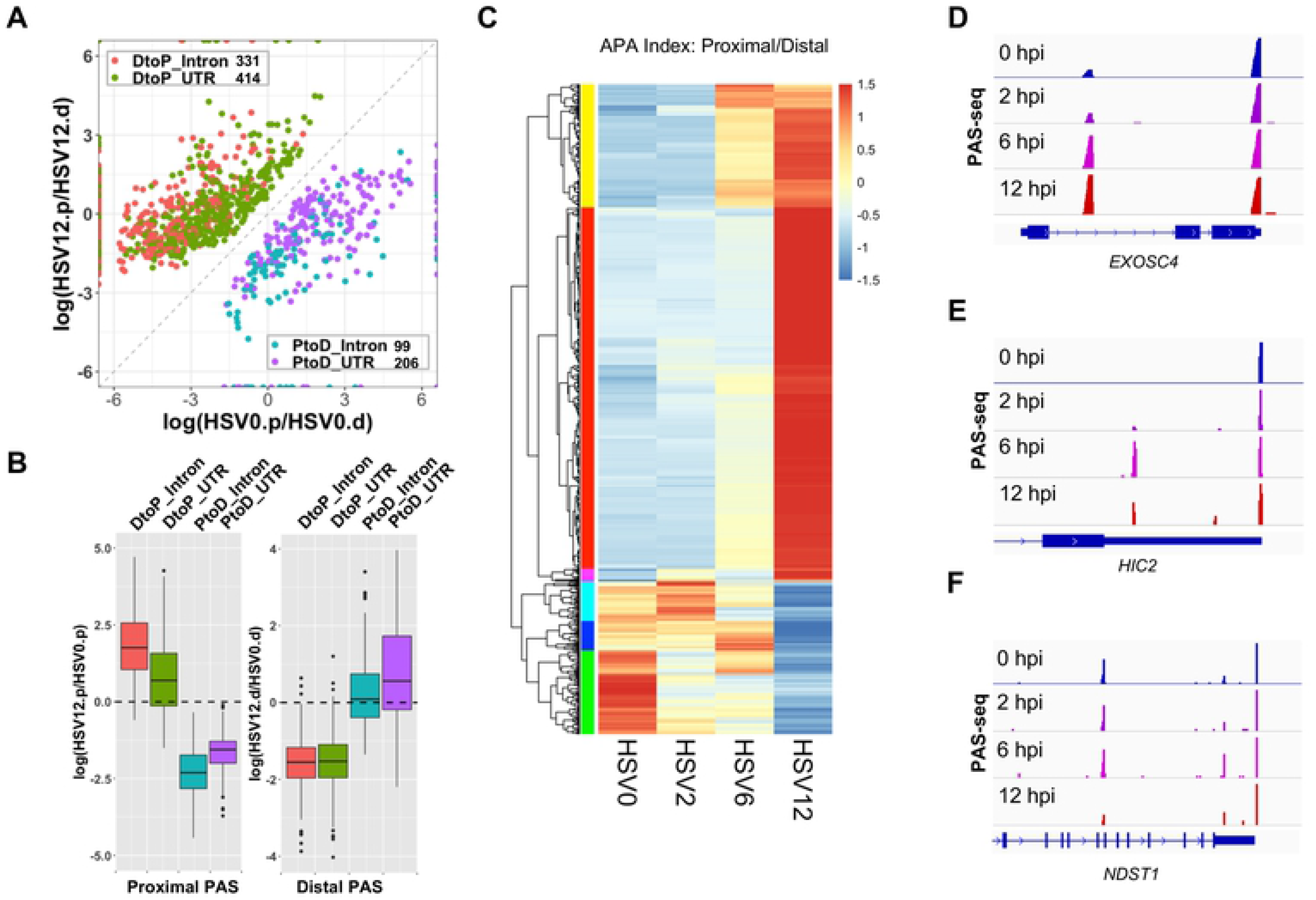
HSV-1 infection induces widespread and dynamic APA changes. (A) A scatter plot of HSV-1-induced significant APA changes (FDR < 0.05, at least 15% of the transcripts shifted). HSV0.p or HSV12.p: read counts for the proximal PAS in cells at 0 or 12 hours post infection by HSV-1; HSV0.d or HSV12.d: read counts for the distal PAS in cells at 0 or 12 hours post infection by HSV-1. DtoP: a distal to proximal shift; PtoD: a proximal to distal shift. Intron: shifts involving an intronic PAS. UTR: both PAS are located in the 3’ UTR. (B) Relative read count changes at proximal (.p) or distal (.d) PAS in the four groups of APA shifts. (C) A heat map showing the APA index (proximal/distal read count ratio) of all the genes with significant APA shifts as shown in (A). Data was scaled by row. Color bars on the left show 6 groups that displayed similar patterns. (D-F) PAS-seq tracks of example genes.

To determine the kinetics of APA changes during HSV-1 infection, we compared the APA index (read count ratio between proximal and distal PAS) of the 1,050 genes (Fig. 1C). The greatest shifts in APA profile occurred between 6 and 12 hpi. However, multiple different kinetic patterns were observed for the timing and magnitude of APA changes (Fig. 1C, see the colored sidebars for classification), indicating that multiple mechanisms are involved in regulating the APA of host genes. Three examples were provided to illustrate the different kinetic groups. For example, polyadenylation of the *EXOSC4* transcripts shifted from a PAS in the 3’ UTR to a proximal intronic PAS (Fig. 1D). The majority of the APA shift occurred between 2 and 6 hpi and a modest further shift was observed between 6 and 12 hpi. Similarly, polyadenylation of *HIC2* transcripts shifted to a proximal intronic PAS between 2 and 6 hpi. However, the usage of this intronic PAS decreased subsequently (Fig. 1E). Finally, a PtoD shift was observed for *NDST1* transcripts and the majority of the APA change occurred between 6 and 12 hpi (Fig. 1F). Together, these data demonstrated that HSV-1 infection induces widespread APA changes, the majority of which shift from distal to proximal PAS. These APA changes follow multiple kinetic patterns, indicating that different mechanisms might be involved in HSV-1-mediated APA regulation of host genes.

### The relationship between the HSV-1-induced APA changes and transcription

HSV-1-induced APA changes could be due to changes in PAS selection during transcription and/or selective loss of individual APA isoforms. To distinguish between these mechanistic models, we directly compared our PAS-seq data with nascent RNA sequencing (4sU-seq) data, which provides information on transcription activities[13]. Meta-analyses of 4sU-seq signals in mock or HSV-1-infected cells along the genes that showed HSV-1-induced higher usage of upstream PAS (DtoP) revealed two interesting differences. First, although the 4sU-seq signals are similar at transcription start sites (TSS), the signal intensities are lower within the gene body in HSV-1-infected cells (Fig. 2A, marked by a red arrow), indicating loss of transcription activity within this region. Second, the 4sU-seq signals downstream of transcription end site (TES) in HSV-1-infected cells are higher than those in mock treated cells, consistent with DoTT (Fig. 2A, marked by a red arrow). To better monitor potential changes in transcription activities near the regulated PAS, we focused on DtoP shifts involving intronic proximal PAS. Importantly, accumulation of 4sU-seq signals were observed at these intronic PAS followed by an abrupt decrease in HSV-1-infected cells (Fig. 2B, a quantitative comparison for individual PAS is shown in Supplemental Fig. 2). This pattern is a hallmark of transcriptional termination [5], suggesting that the observed higher PAS-seq signals at these intronic PAS are, at least in part, due to higher usage of these PAS during transcription. This is exemplified for the gene *TOB2* (Fig. 2C). Here, an intronic PAS in *TOB2* is activated in the HSV-1-infected cells. Concomitantly, 4sU-seq signals accumulated in this region followed by a decrease in HSV-1-infected cells, consistent with transcriptional termination. Significantly higher 4sU-seq signals were also observed downstream of the TES consistent with impaired PAS usage and read-through transcription at the canonical downstream PAS (Fig. 2C). As a comparison, we also plotted the 4sU-seq signals in mock and HSV-1-infected cells for genes that displayed PtoD APA changes. Different from the DtoP genes (Fig. 2A-B), the 4sU-seq signals from mock and HSV-1-infected cells were very similar over the gene body for PtoD genes (Fig. 2D), indicating that there was no early transcription termination. Similar to the DtoP genes, however, 4sU-seq signals also accumulated downstream of TES in PtoD genes (Fig. 2D, marked by a red arrow), suggesting that DoTT is a common feature for both classes of genes. When examining the 4sU-seq signals at the intronic PAS of PtoD genes, we observed the opposite pattern than the DtoP genes. A peak followed by a valley pattern was detected at these intronic PAS in mock treated cells (Fig. 2E), indicating polyadenylation at these sites is accompanied by transcriptional termination. By contrast, 4sU-seq signals in HSV-1-infected cells were relatively flat at these intronic PAS, indicating transcriptional readthrough (Fig. 2E, red line). This pattern is exemplified by the *DNAJB6* gene (Fig. 2F). Both polyadenylation of *DNAJB6* transcripts and transcription termination occurred primarily at an intronic PAS in mock treated cells (Fig. 2F). In HSV-1 infected cells, polyadenylation shifted to the distal PAS in the terminal exon, which was accompanied by transcription readthrough at the intronic PAS (Fig. 2F). Together, these data suggest that the HSV-1-induced changes in APA profiles are, at least in part, caused by changes in PAS usage during transcription.

**Figure 2.**
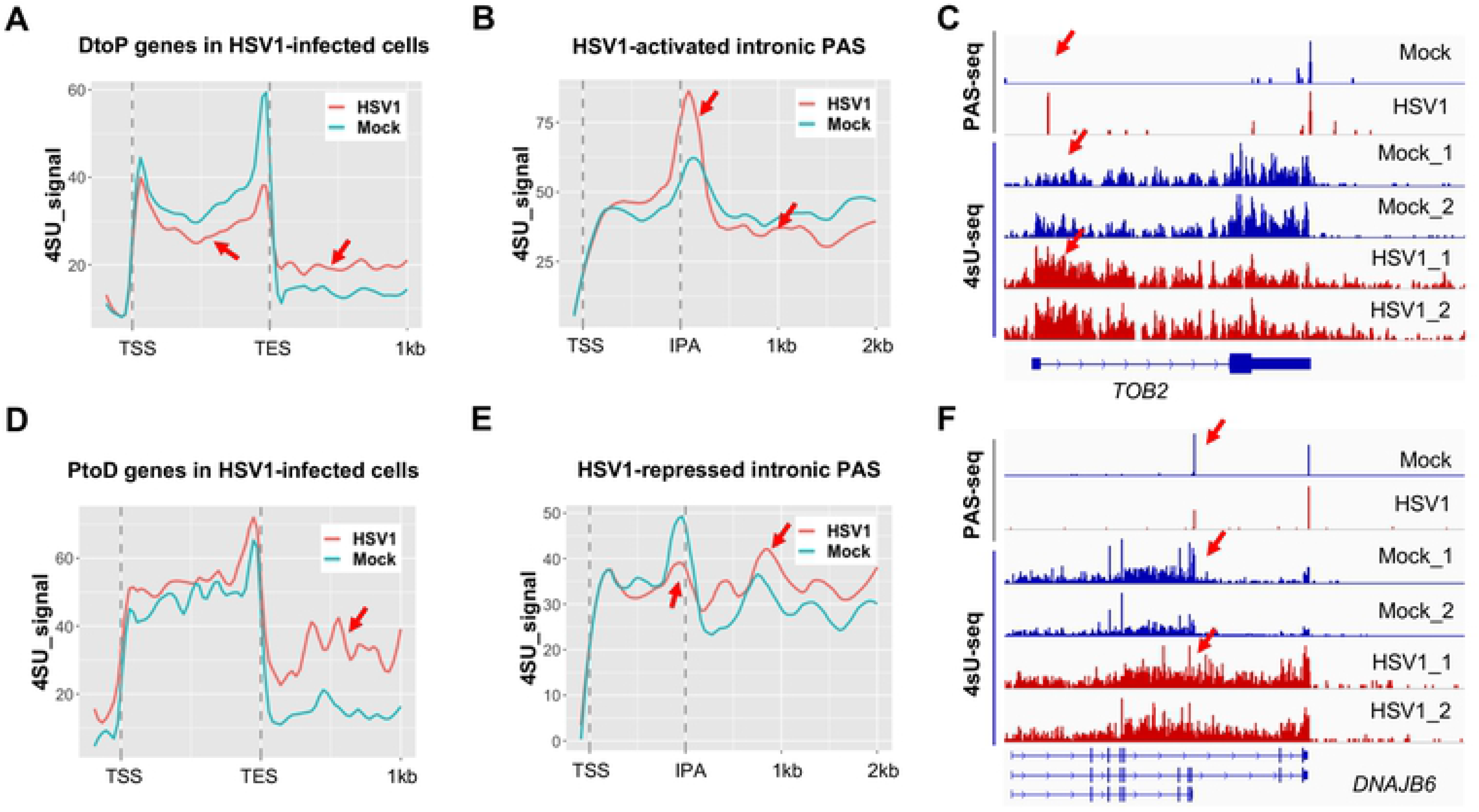
HSV-1-induced APA changes occur, at least in part, co-transcriptionally. (A) 4sU-seq signals at genes that displayed DotP APA shifts in HSV-1 infected cells. Red arrows point out the regions with differences. TSS: transcription start site. TES: transcription end site. (B) 4sU-seq signals at genes that displayed DotP APA shifts in HSV-1 infected cells involving intronic PAS. IPA: intronic polyadenylation site. (C) PAS-seq and 4sU-seq tracks of *TOB2*. Red arrows point to the IPA region. (D) 4sU-seq signals at genes that displayed PtoD APA shifts in HSV-1 infected cells. (E) 4sU-seq signals at genes that displayed PtoD APA shifts in HSV-1 infected cells involving intronic PAS. (F) PAS-seq and 4sU-seq tracks of *DNAJB6*. Red arrows point to the IPA region.

### ICP27-dependent and -independent APA changes during HSV-1 infection

We recently showed that the HSV-1 immediate early factor ICP27 directly interacts with the mRNA 3’ processing factor CPSF and blocks mRNA 3’ end formation [13]. This suggests that ICP27 could be directly involved in HSV-1-mediated APA regulation. To test this possibility, we first compared the APA changes induced by the wild-type and the ΔICP27 HSV-1, in which the ICP27 gene was replaced by lacZ [23], by PAS-seq. The majority of HSV-1-induced APA changes were abolished or diminished in ΔICP27 infected cells (Fig. 3A, compare HSV1 and ΔICP27), strongly suggesting that ICP27 is required for HSV-1-mediated APA regulation. Interestingly, however, when comparing the APA profiles of mock and ΔICP27 infected cells, we detected 1,435 significant APA changes and the majority of these APA changes (1,109 genes or 77%) are DtoP shifts (Fig. 3B). Therefore, the ΔICP27 virus induced an even greater number of APA changes than the wild-type virus. The proximal PAS involved in ΔICP27- and wild-type HSV-1-induced APA changes are largely distinct with relatively small overlap (310 genes in the overlap, Fig. 3C). An overall comparison of mock, HSV-1, and ΔICP27 at all PAS that displayed significantly different usage in either wild-type or ΔICP27 HSV-1-infected cells is shown in Supplemental Fig. 3. These data demonstrate that HSV-1 can induce both ICP27-dependent and -independent APA changes. For example, HSV-1 infection activated an intronic PAS in *EXOSC4* transcripts, and a similar activation was not observed in ΔICP27-infected cells (Fig. 3D). By contrast, an intronic PAS in *CHTOP* is ICP27-independent as it was similarly activated in both wild-type and ΔICP27 HSV-1-infected cells (Fig. 3E). Finally, an intronic PAS in *GLIS2* was only activated in ΔICP27-but not in wild-type HSV-1-infected cells (Fig. 3F). Therefore, these results suggest that HSV-1 can induce APA changes through multiple mechanisms.

**Figure 3.**
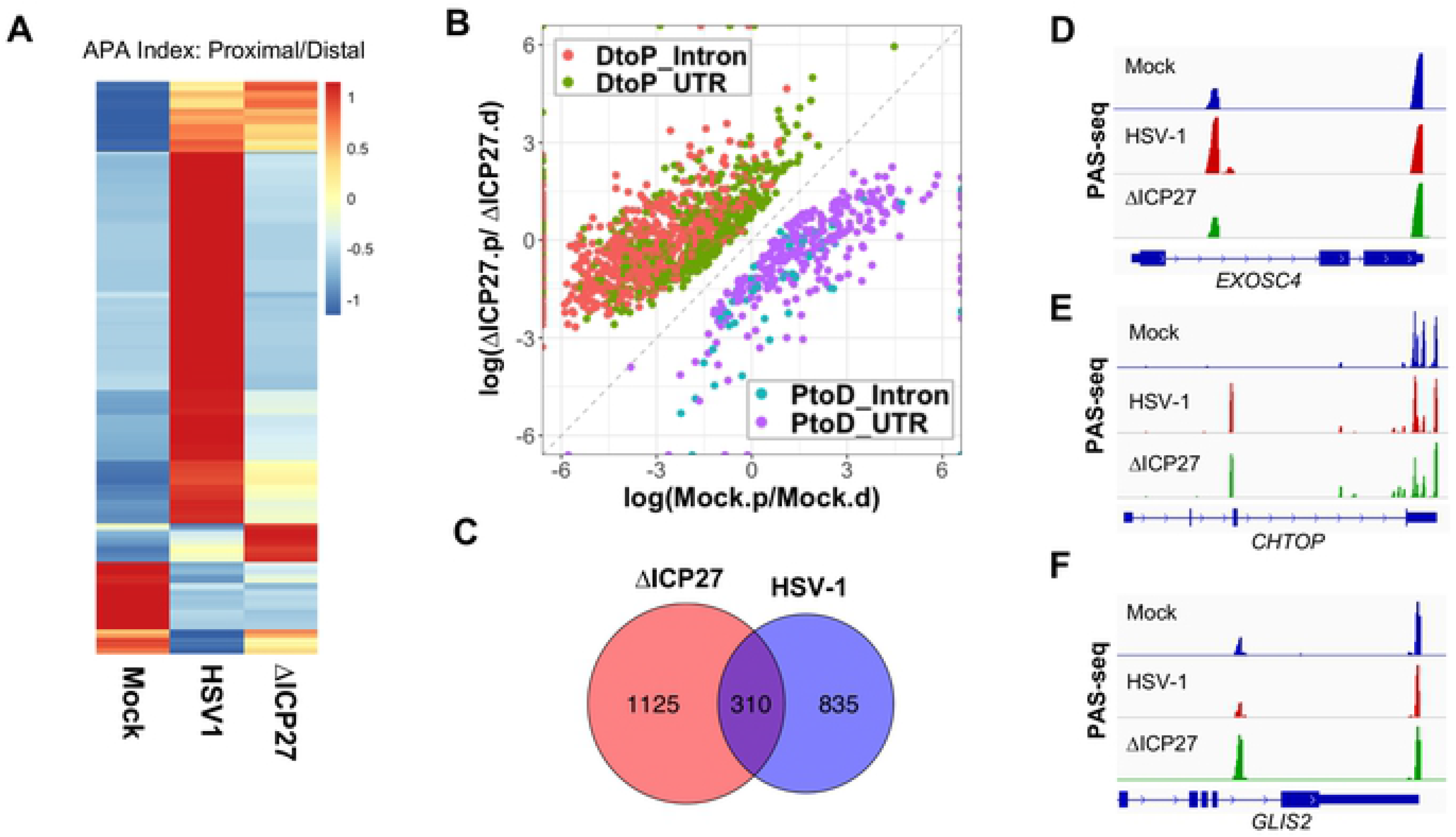
ICP27-dependent and -independent HSV-1-induced APA changes. (A) A heat map showing the APA index of all the genes that displayed significant APA changes as shown in Fig. 1A in mock, wild-type or ΔICP27 HSV-1-infected cells. Data was scaled by row. (B) A scatter plot showing significant APA differences between mock- and ΔICP27 HSV-1-infected cells. Similar to Fig. 1A. (C) A Venn diagram showing the overlap between proximal PAS that displayed significant APA changes induced by the wild-type or ΔICP27 HSV-1. (D-F) PAS-seq tracks of mock-, wild-type HSV-1, or ΔICP27 HSV-1-infected cells at 8 hpi.

### Mechanisms for HSV-1-mediated APA regulation

Our data suggest that ICP27 is necessary for the majority of APA changes induced by wildtype HSV-1 infection. We thus wanted to determine if ICP27 is sufficient to regulate APA. Based on RNA-seq analyses, the Krause laboratory recently provided evidence that ectopically expressed ICP27 regulates APA [24]. However, RNA-seq is not ideal for APA analysis as it lacks the sensitivity to detect APA changes of modest magnitude or those involving closely located alternative PAS [8]. To overcome these limitations, we performed PAS-seq analysis of mock-transfected or ICP27 over-expressing cells. Overexpression of ICP27 induced significant APA changes in 169 genes, the vast majority of which (154 genes or 91%) were DtoP shifts (Fig. 4A). Among these DtoP shifts, 111 genes or 72% shifted to a proximal intronic PAS (DtoP_Intron, Fig. 4A). The majority of ICP27 overexpression-induced APA changes were also induced by HSV-1 infection (p < 5.1e-71, hypergeometric test; Supplementary Fig. 4A). However, the number of genes regulated by ICP27 overexpression was significantly lower compared to HSV-1-regulated APA events. Thus, although ICP27 is necessary for a majority of HSV1-induced APA changes, it is not sufficient to induce these changes.

**Figure 4.**
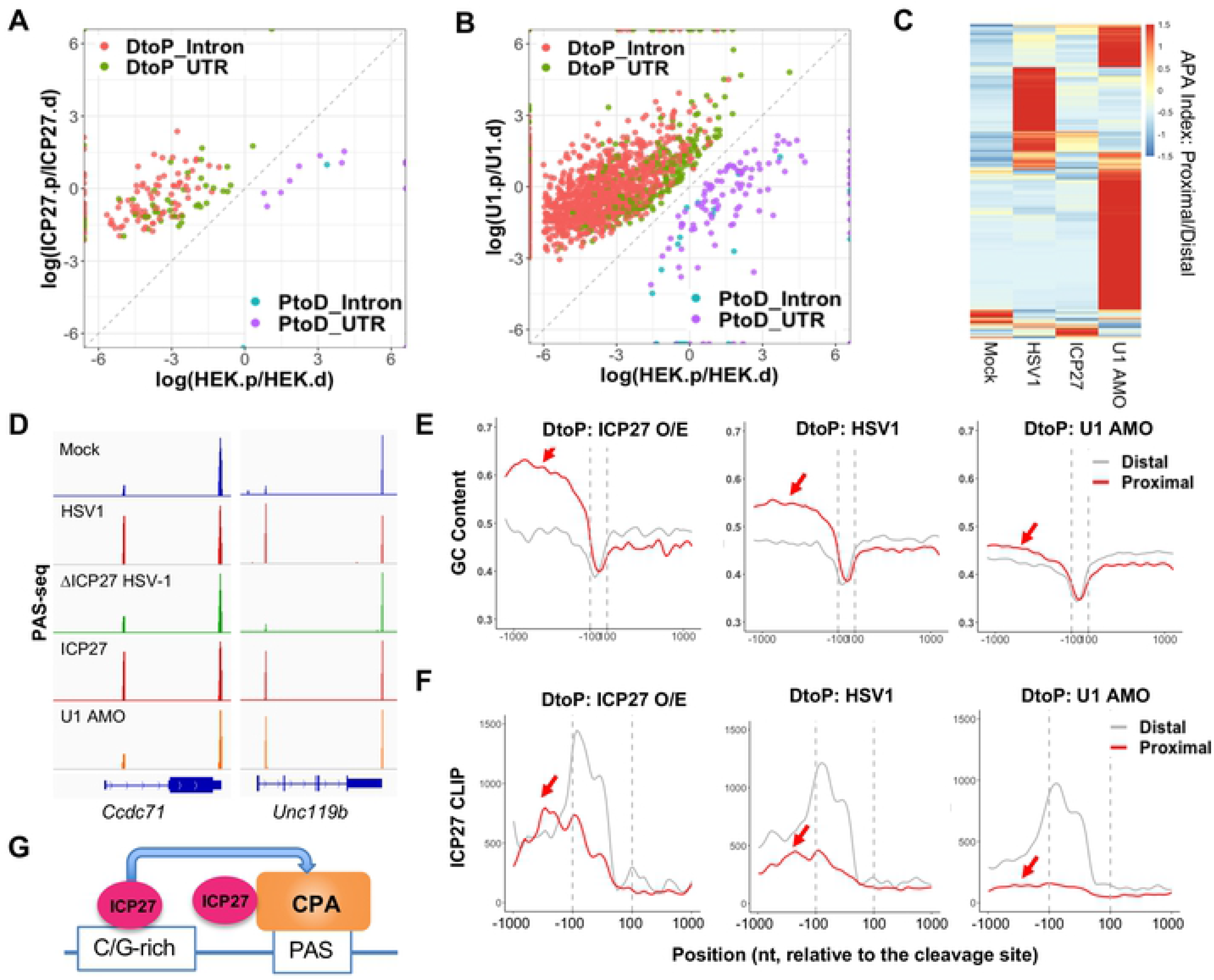
Mechanism of HSV-1-mediated APA regulation. (A) A scatter plot showing the significant APA changes induced by ICP27 over-expression in HEK293T cells. Color scheme and labeling are similar to Fig. 1A. (B) A scatter plot showing the significant APA changes induced by U1 antisense morpholino oligo (AMO). (C) A heat map of APA index of all genes that displayed significant APA changes induced by HSV-1-infection, ICP27 over-expression, or U1 AMO. Data are scaled by row. (D) PAS-seq tracks for two example genes. (E) GC content at the proximal and distal PAS of genes that displayed significant DtoP shifts induced by ICP27 over-expression (O/E), HSV-1 infection (HSV1), or U1 AMO treatment. (F) ICP27 CLIP signals at the proximal and distal PAS of genes that displayed significant DtoP shifts induced by ICP27 over-expression (O/E), HSV-1 infection (HSV1), or U1 AMO treatment.

Cleavage and polyadenylation at intronic PAS are generally inhibited by the U1 snRNP [12]. As ICP27 overexpression primarily activates intronic PAS, it was proposed that ICP27 may modulate APA by blocking U1 snRNP activity [24]. To test this model, we transfected U1 antisense morpholino oligo (AMO) into cells, which blocks U1 snRNA-RNA interactions and thereby inhibiting U1 activity. PAS-seq analysis showed that U1 AMO treatment resulted in significant APA changes in 1,999 genes, the majority of which (1,867 genes or 93%) were DtoP shifts (Fig. 4B). Consistent with previous studies, the majority of these APA changes (1,646 genes or 82% of the total) involve the activation of an intronic PAS (Fig. 4B). A comparison between U1 AMO-, HSV-1- and ICP27-induced APA changes revealed largely distinct patterns with small overlaps (Fig. 4C). For example, among the 169 genes whose APA is regulated by ICP27, only 32 (19%) are also regulated by U1 snRNP (Supplemental Fig. 4B). Similarly, 13% of HSV1-induced APA changes were also induced by U1 AMO (Supplemental Fig. 4C). For examples, ICP27 is necessary and sufficient to activate an intronic PAS in an intronic PAS in *CCDC71* and *UNC119b* (Fig. 4D). However, the intronic PAS of *UNC119b*, but not *CCDC71,* was induced by U1 AMO treatment (Fig. 4D). These data suggest that HSV-1-mediated APA regulation does not involve a general inhibition of U1 snRNP.

To begin to understand the molecular basis for the specificity of HSV-1- and ICP27-induced activation of intronic PAS, we examined the sequences of the regulated PAS. Comparison of the intronic PAS activated by either ICP27 or HSV-1 with the corresponding distal PAS in the 3’ UTR, revealed a higher GC content at the intronic PAS (Fig. 4E). ICP27-activated intronic proximal PAS are highly G/C-rich in the region upstream of cleavage sites (Fig. 4E, left panel). HSV-1-activated intronic proximal PAS show an intermediate GC content (Fig. 4E, middle panel). By contrast, intronic PAS activated by U1 suppression had a lower GC content (less than 50%, Fig. 4E, right panel). We have previously shown that ICP27, when bound to upstream GC-rich sequences, can activate PAS (Fig. 4G) [13]. To test if the GC contents of the different classes of intronic PAS impact ICP27 binding, we took advantage of the ICP27 CLIP-seq dataset that we generated previously [13]. Interestingly, we commonly observed high ICP27 CLIP-seq signals upstream of ICP27-activated intronic PAS and intermediate levels of ICP27 CLIP-seq signals at HSV-1-induced intronic PAS (Fig. 4F, left and middle panels). By contrast, very little ICP27 binding was detected upstream of U1-regulated intronic PAS (Fig. 4F, right panel). Thus, the ICP27 CLIP-seq signal intensities at these intronic PAS are highly consistent with the respective GC content. These observations are consistent with the model that ICP27 activates specific PAS by binding to GC-rich upstream sequences during HSV-1 infection, and that HSV-1-mediated APA regulation does not involve a general inhibition of U1 snRNP.

### Export of HSV-1-induced APA isoforms

Finally, we wanted to determine how HSV-1-induced APA changes regulate the export of the corresponding transcripts. To address this question, we analyzed a RNA-seq dataset that we have recently generated for chromatin, nucleoplasmic, and cytoplasmic fractions of mock- or HSV-1-infected cells at 8 hpi [17]. Nuclear RNAs, such as scaRNA2, which functions in snRNA modification in the nuclear Cajal bodies [25], were highly enriched in the chromatin and nucleoplasmic fractions (Supplemental Fig. 5A). By contrast, RNAs of highly expressed house-keeping genes were primarily detected in the cytoplasm (Supplemental Fig. 5B), suggesting that fractionation was successful. To measure the overall export efficiencies of HSV-1 target APA genes, we performed a meta-analysis of all 1,050 genes that displayed significant APA changes in HSV-1-infected cells. As shown in Fig. 5A, in the chromatin fraction of mock treated cells, RNA-seq signals were prominent in both gene body and the downstream region, the latter of which most likely resulted from uncleaved precursor mRNA. This is consistent with the genome-wide trend [17]. By contrast, both nuclear and cytoplasmic fractions showed heightened signals in the gene body and lower signal in the region downstream of TES (Fig. 5A). These results demonstrate that, in mock treated cells, transcripts are efficiently cleaved and the mature mRNAs are released into the nucleoplasm and subsequently exported into the cytoplasm. By sharp contrast, in the chromatin fraction of HSV-1-infected cells, RNA-seq signals in the region downstream of TES were significantly elevated, consistent with the known transcription termination defect [14,17] (Fig. 5B). Higher RNA-seq signals downstream of the TES were also observed in the nucleoplasmic, but not cytoplasmic fraction (Fig. 5B), indicating that the transcripts that extended past the TES were released into the nucleoplasm, but not exported. The release of DoTT transcripts could be due to cleavage/polyadenylation downstream of the normal TES. Indeed, as shown in Fig. 5C, elevated levels of PAS-seq signals were observed downstream of *HNRNPA2B1* TES (PAS-seq tracks, peaks downstream of the TES are marked by a red arrow). On the global level, we compared the PAS-seq reads in the 5 kilobase (kb) region downstream of the annotated TES for all PtoD genes and found that, indeed, there are significantly higher PAS-seq reads within this region in HSV-1-infected cells compared to mock treated cells (Fig. 5D, p=0.002, Wilcoxon test). The PAS-seq reads in the downstream regions contain the canonical poly(A) signal, AWTAAA hexamer, at −20 nt position (Supplementary Fig. 6A), and do not have a poly(A) run downstream of the cleavage sites (Supplementary Fig. 6B), strongly suggesting that these PAS-seq reads are due to the usage of cryptic PAS and not due to a potential technical artifact such as internal priming. These data suggest that HSV-1-induced extended transcripts as a result of DoTT are released into the nucleoplasm by cleavage/polyadenylation at cryptic PAS in the downstream region. However, these transcripts are not efficiently exported into the cytoplasm (Fig. 5B-C).

**Figure 5.**
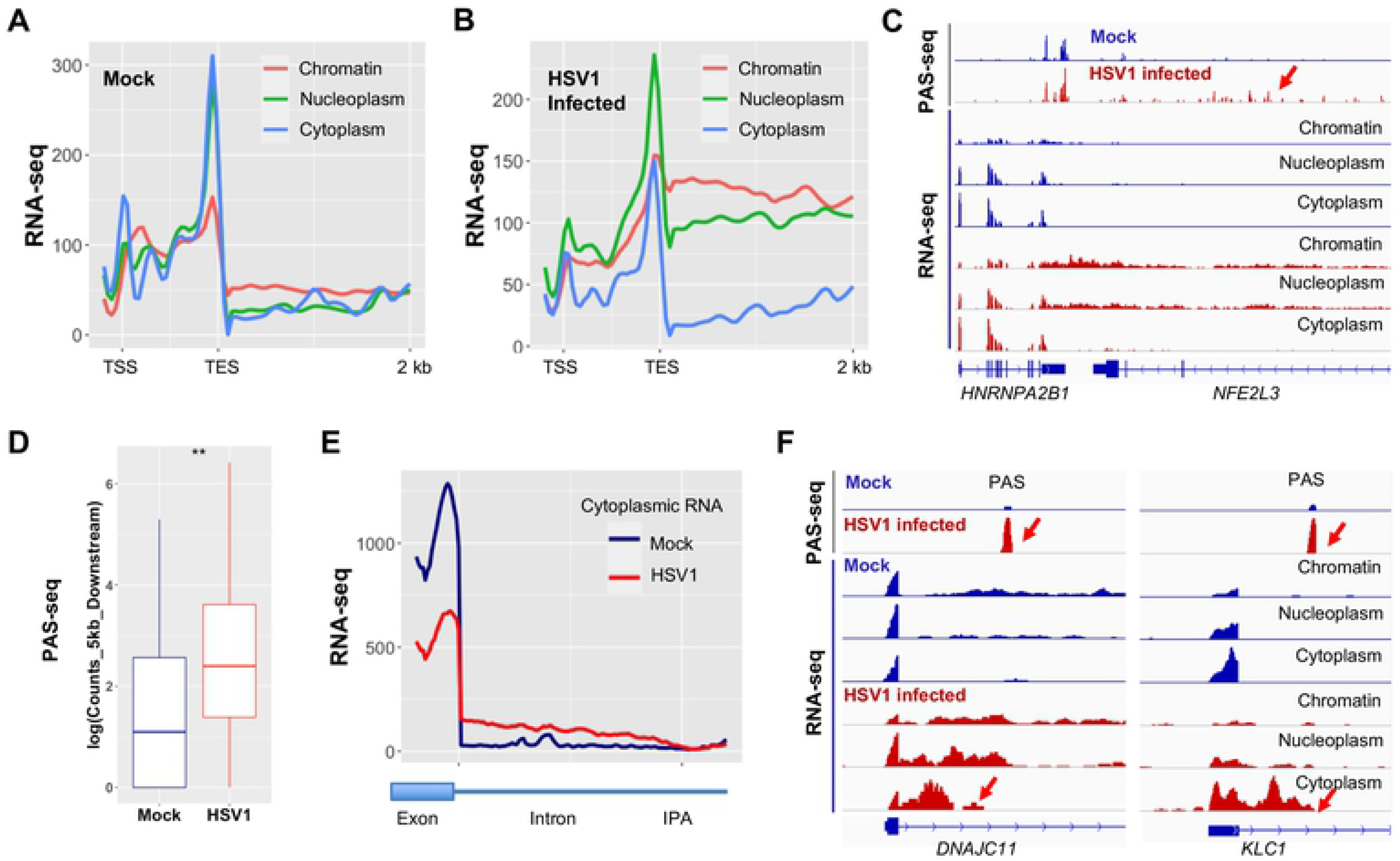
HSV-1-mediated APA regulation and mRNA export. Average RNA-seq signals for genes that display significant HSV-1-induced APA changes in chromatin, nucleoplasm, and cytoplasm fractions in mock-infected (A) or HSV-1-infected cells (B). (C) PAS-seq and RNA-seq tracks for HNRNPA2B1 gene. Red arrow points to the PAS-seq signals downstream of the normal TES. (D) PAS-seq signals in the 5kb region downstream of the normal TES in mock- and HSV-1-infected cells. ** p value=0.0039. (E) Cytoplasmic RNA-seq signals for HSV-1-induced DtoP_intron APA changes. IPA: intronic poly(A) site. (F) PAS-seq and RNA-seq tracks for two example genes. Red arrows point to the IPA.

HSV-1 infection activates intronic PAS in a large number of genes (Fig. 1A). The resultant transcripts are predicted to encode aberrant truncated proteins. To monitor the fate of these RNAs, we performed a meta-analysis of the region from the upstream exon to the intronic PAS for DtoP_intronic genes, which distinguishes the spliced and polyadenylated APA isoforms (Fig. 5E). Signals from the upstream exon reflect both spliced and polyadenylated transcripts whereas the signals in the intronic region are only derived from the unspliced polyadenylated isoform. In the cytoplasm of mock infected cells, high RNA-seq signals were observed for the upstream exon while almost no signals were detected in the intronic regions (Fig. 5E, blue line), suggesting that only fully spliced transcripts are exported. However, RNA-seq signals decreased in the upstream exon region, but accumulated between the upstream exon and the intronic PAS in the cytoplasm of HSV-1-infected cells (Fig. 5E, red line). This suggests that the transcripts polyadenylated at intronic PAS are exported into the cytoplasm. Two examples were provided in Fig. 5F. For both *DNAJC11* and *KLC1*, their transcripts are efficiently spliced in mock treated cells, but HSV-1 infection activates a PAS within the first intron, as shown by the PAS-seq data (Fig. 5F, PAS-seq tracks, activated intronic PAS are marked by red arrows). Our fractionation RNA-seq data showed that these truncated RNA isoforms are exported into the cytoplasm (Fig. 5F, RNA-seq tracks, cytoplasmic tracks are marked by red arrows). Based on these observations, we conclude that the transcripts of the APA target genes are exported less efficiently and that the truncated transcripts polyadenylated at intronic PAS are exported. We have attempted to determine if the exported intronically polyadenylated APA isoforms are translated by using the Ribo-seq dataset that we generated previously [14]. However, the Ribo-seq signal density was not sufficient for reliable analyses and, as a result, it remains unclear whether these HSV-1-induced shorter APA isoforms are translated.

## Discussion

### Mechanism and functional impact

mRNA 3’ end processing and transcription termination are tightly coupled processes. Viral infections (HSV-1 and IAV) and cellular stresses (salt/osmotic stress and heat shock) induce DoTT/DoG [14–16]. Meanwhile, several pathogens, including viruses (HCMV and VSV) and bacteria (listeria and salmonella), as well as arsenic stress causes widespread APA changes [18–20,26]. However, no study has characterized these two processes in response to the same pathogen or stress. In this report, we performed extensive transcriptomic analyses of wild-type and mutant HSV-1-infected cells and found that lytic HSV-1 infection induced widespread APA changes in host transcripts, the majority of which shifted to upstream PAS. HSV-1-mediated APA regulation requires the viral immediate early factor ICP27 as well as other viral factors, but does not involve a general inhibition of U1 snRNP. Interestingly, HSV-1 induces both activation of upstream PAS with pre-mature transcription termination and a termination defect. Activation of upstream intronic PAS produces truncated transcripts that are exported into the cytoplasm. By contrast, although extended transcripts due to DoTT can be cleaved and polyadenylated at downstream cryptic PAS, these transcripts are sequestered in the nucleus. Together, these results demonstrate that HSV-1-mediated regulation of APA and transcription termination profoundly reprograms host transcriptomes (Fig. 6).

**Figure 6.**
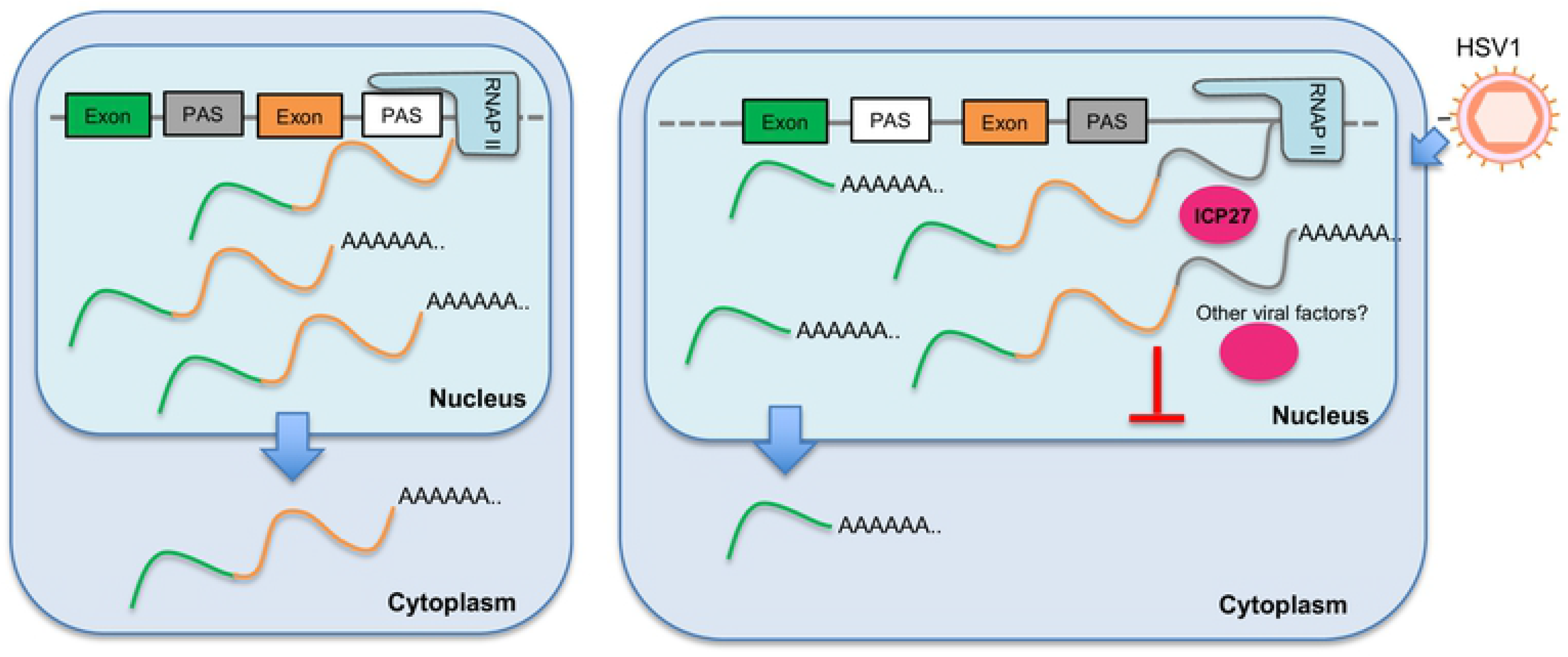
A model for HSV-1-mediated APA regulation. Please see text for details.

Although widespread APA changes have been described for a number of pathogen-infected cells and for cells exposed to arsenic stress [18–20,26], the underlying mechanism remains poorly understood. Our data suggest that the viral immediate early factor ICP27 contributes to HSV-1-induced APA changes. We have previously shown that ICP27 has bimodal activities: it broadly inhibits mRNA 3’ processing through direct interactions with the 3’ processing factor CPSF, but can activate PAS that contain GC-rich upstream sequences [13]. Indeed, both HSV-1 infection and over-expression of ICP27 can activate upstream intronic PAS and these PAS contains GC-rich upstream sequences (Fig. 4E-F). 3’ processing at these intronic PAS induces early termination (Fig. 2A-C). On the other hand, the corresponding downstream PAS in these genes lack GC-rich upstream sequences and are thus inhibited, leading to DoTT at these sites. Thus, the bimodal activities of ICP27 provide an explanation for the paradoxical observation of early termination and DoTT in these genes. On the other hand, the ΔICP27 virus still induce a large number of APA, suggesting that other factors are also involved. Similarly, we have previously shown that multiple viral factors contribute to HSV-1-induced DoTT [13]. Since multiple pathogens and stress induce similar changes in both APA and DoTT, it is likely that a common mechanism underlies these phenomena. One possibility is that viral infections and cellular stress may alter the activity of RNAPII. In addition to its role in transcribing genes, RNAPII also plays an essential role in coordinating transcription and RNA processing primarily through its C-terminal domain. Both phosphorylation and dephosphorylation of RNAPII CTD have been shown to influence termination [4–6,27]. For example, pharmacological or genetic inhibition of Cdk12, which phosphorylates RNAPII CTD at serine 2, leads to activation of intronic PAS and premature termination [28,29]. On the other hand, PP1 or PP2A, phosphatases that dephosphorylate RNAPII CTD, play essential roles in regulating transcription pausing and termination [5,30–32]. Previous studies provided evidence that HSV-1 infection induces aberrant CTD phosphorylation and partial degradation of RNAPII [33][34]. It will be important to characterize RNAPII post-translational modifications and interactomes in pathogen-infected and in stressed cells and determine if/how such changes contribute to the virus-induced APA changes and DoTT.

The functional consequence of pathogen/stress-induced DoTT and APA changes remains unclear. The most important functions of stress responses are to: 1) shut down the expression of most genes to avoid accumulation of aberrant proteins; 2) activate stress response genes to stabilize and repair biomolecules [35]. Similarly, when a pathogen infects a host cell, it shuts down host gene expression and hijacks the host machinery to express genes of the pathogen. Both DoTT and APA changes could contribute to the repression of cellular genes. DoTT interferes with the transcription cycle and prevents mRNA biogenesis. Consistent with previous reports [17], our results showed that at least some of the read-through transcripts as a result of DoTT are in fact cleaved and polyadenylated, and released into the nucleoplasm (Fig. 5). However, they are not efficiently exported. On the other hand, HSV-1-induced activation of upstream intronic PAS leads to the production of truncated transcripts that can be exported (Fig. 5E). However, these transcripts do not encode the functional full-length proteins. Therefore, both DoTT and APA may function in host shutoff (for pathogens) or repressing bulk gene expression (for stresses). Alternatively, the DoTT and APA changes observed in pathogen-infected or stressed cells could represent a host defense mechanism. Previous studies provided evidence that arsenic stress-induced APA isoforms with shorter 3’ UTRs, which can evade RNA degradation, are thus better preserved [20]. This may facilitate better recovery from stress. VSV-induced APA changes have been shown to modulate the innate immunity response [19]. In summary, pathogen- and stress-induced APA changes may function in host shut-off or in host defense, and these two mechanisms are not mutually exclusive.

## Acknowledgement

We thank the UCI GHTF for sequencing. This study was supported by the following grants: NIH GM090056 and GM128441 to Y.S., ERC Consolidator award (ERC-2016-CoG 721016-HERPES) and DFG grants DO1275/6-1 to L.D. and FR2938/9-1 to C.C.F. A.W.W. was the recipient of a generous grant from the Alexander von Humboldt Foundation.

## Methods

### 1. Cell Culture, viruses and infection

HEK293 and HeLa cell lines were cultured in Dulbecco’s modified Eagle medium (DMEM) with 10% fetal bovine serum (FBS). All cells were incubated at 37°C in a 5% (v/v) CO2-enriched incubator. Virus stocks for wild-type HSV-1 strain KOS as well as the ICP27 null mutant (strain KOS) [23] were produced on complementing Vero 2-2 cells [36]. Cells were infected with an MOI of 10 unless otherwise specified and incubated at 37°C until cells were harvested at the specified time points. For anti-sense morpholino oligo treatment, HEK293 cells were treated with 50 μM U1 antisense morpholino oligo (AMO) (Gene tools) and 10 μM Endo-Porter (Gene tools). After 48 hours, RNA was extracted by using Trizol (Ambion).

### 2. PAS-seq

Total RNA was extracted with Trizol as per manual (Life technologies), 10 μg total RNA was fragmented with fragmentation reagent (Ambion) at 70 °C for 10 minutes followed by precipitation with ethanol. After centrifugation, RNA was dissolved and Reverse transcription was performed with PASSEQ7-2 RT oligo: [phos]NNNNAGATCGGAAGAGCGTCGTGTTCGGATCCATTAGGATCCGAGACGTGTGCTCTTCCGATCTTTTTTTTTTTTTTTTTTTT[V-Q] and Superscript III. cDNA was recovered by ethanol precipitation and centrifugation. 120-200 nucleotides of cDNA was gel-purified and eluted from 8% Urea-PAGE. Recovered cDNA was circularized with Circligase™ II (Epicentre) at 60 °C overnight. Buffer E (Promega) was added in cDNA and heated at 95 °C for 2 minutes, and then cool to 37 °C slowly. Circularized cDNA was linearized by adding BamH I (Promega). cDNA was centrifugated after ethanol precipitation. PCR was carried out with primers PE1.0 and PE2.0 containing index. Around 200 bp of PCR products was gel-purified and submitted for sequencing (single read 100 nucleotides).

### 3. PAS-Seq Data Analysis

From the raw PAS-seq reads, first those with no poly(A) tail (less than 15 consecutive “A”s) were filtered out. The rest were trimmed and mapped to hg19 genome using STAR. If 6 consecutive “A”s or more than 7 “A”s were observed in the 10 nucleotide downstream of PAS for a reported alignment, it was marked as a possible internal priming event and removed. The bigwig files were then generated for the remaining reads using deepTools (v2.4) with “normalizeUsingRPKM” and “ignoreDuplicates” parameters [37].

Next, the locations of 3’ ends of the aligned reads were extracted and those in 40nt of each other were merged into one to provide a list of potential PAS for human. This list was then annotated based on the canonical transcripts for known genes. The final read count table was created using the reads with their 3’ ends in −40nt to 40nt of these potential PAS.

Alternatively polyadenylated PAS in different experimental conditions were identified using diffSpliceDGE and topSpliceDGE from edgeR package(v3.8.5) [38]. This pipeline first models the PAS read counts for all PAS, then compares the log fold change of each PAS to the log fold change of the entire gene. This way, these functions, primarily used to find differential exon usage, generate a list of sites with significant difference between our PAS-seq samples. From this list, those with a FDR value less than 0.05 and more than 15% difference in the ratio of PAS read counts to gene read counts (normalized by sequencing depth) between samples were kept, and finally for each gene the top two were chosen based on P-value and marked distal or proximal based on their relative location on the gene. For PAS-seq comparisons without replicates, Fisher’s exact test was used to compare read counts at a PAS and the total read counts from the same gene. The P values were adjusted by the Benjamini–Hochberg method for calculating the FDR.

For the genes with alternatively polyadenylated sites (target genes), the log2 of ratio of read counts in the distal site to the read counts in the proximal site was calculated and illustrated as a heatmap in Fig. 1C, 2A, and 3C with pheatmap in R. The heatmap is hierarchically clustered using Pearson correlation of the gene profiles in different experiments.

### 4. General Analysis

The computational analyses and visualization if not specified otherwise, were done in Python 2.7. Where necessary, conversion between BAM and BED files were done using BEDTools (v2.25.0) and BAM files were sorted or indexed via SAMtools (v1.1) [39][40]. Meta-analyses of read distribution were performed using deeptools [37].

### Data and Software Availability

RNA-seq data on the subcellular RNA fractions and 4sU-seq data were previously published [13,17]. PAS-seq data have been deposited to the GEO database (GSE151104).

## Reference

1. Colgan DF, Manley JL. Mechanism and regulation of mRNA polyadenylation. Genes Dev. 1997;11: 2755–2766. Available: http://www.ncbi.nlm.nih.gov/entrez/query.fcgi?cmd=Retrievedb=PubMeddopt=Citationlist_uids=9353246

2. Zhao J, Hyman L, Moore C. Formation of mRNA 3’ ends in eukaryotes: mechanism, regulation, and interrelationships with other steps in mRNA synthesis. Microbiol Mol Biol Rev. 1999/06/05. 1999;63: 405–445. Available: http://www.ncbi.nlm.nih.gov/pubmed/10357856

3. Chan S, Choi EA, Shi Y. Pre-mRNA 3’-end processing complex assembly and function. Wiley Interdiscip Rev RNA. 2011/10/01. 2011;2: 321–335. doi:10.1002/wrna.54

4. Richard P, Manley JL. Transcription termination by nuclear RNA polymerases. Genes Dev. 2009/06/03. 2009;23: 1247–1269. doi:10.1101/gad.1792809

5. Eaton JD, West S. Termination of Transcription by RNA Polymerase II: BOOM! Trends in Genetics. Elsevier Ltd; 2020. pp. 664–675. doi:10.1016/j.tig.2020.05.008

6. Proudfoot NJ. Transcriptional termination in mammals: Stopping the RNA polymerase II juggernaut. Science (80-). 2016;352: aad9926–aad9926. doi:10.1126/science.aad9926

7. Tian B, Manley JL. Alternative polyadenylation of mRNA precursors. Nat Rev Mol Cell Biol. 2016;18: 18–30. doi:10.1038/nrm.2016.116

8. Shi Y. Alternative polyadenylation: new insights from global analyses. RNA. 2012/10/26. 2012;18: 2105–2117. doi:10.1261/rna.035899.112

9. Mayr C. Evolution and Biological Roles of Alternative 3′UTRs. Trends Cell Biol. 2016;26: 227–237. doi:10.1016/J.TCB.2015.10.012

10. Di Giammartino DC, Nishida K, Manley JL. Mechanisms and consequences of alternative polyadenylation. Mol Cell. 2011/09/20. 2011;43: 853–866. doi:10.1016/j.molcel.2011.08.017

11. Shi Y, Manley JLJL. The end of the message: multiple protein--RNA interactions define the mRNA polyadenylation site. Genes Dev. 2015;29: 889–897. doi:10.1101/gad.261974.115

12. Kaida D, Berg MG, Younis I, Kasim M, Singh LN, Wan L, et al. U1 snRNP protects pre-mRNAs from premature cleavage and polyadenylation. Nature. 2010/10/01. 2010;468: 664–668. doi:10.1038/nature09479

13. Wang X, Hennig T, Whisnant AW, Erhard F, Prusty BK, Friedel CC, et al. Herpes simplex virus blocks host transcription termination via the bimodal activities of ICP27. Nat Commun. 2020;11. doi:10.1038/s41467-019-14109-x

14. Rutkowski AJ, Erhard F, L’Hernault A, Bonfert T, Schilhabel M, Crump C, et al. Widespread disruption of host transcription termination in HSV-1 infection. Nat Commun. 2015;6. doi:10.1038/ncomms8126

15. Zhao N, Sebastiano V, Moshkina N, Mena N, Hultquist J, Jimenez-Morales D, et al. Influenza virus infection causes global RNAPII termination defects. Nat Struct Mol Biol. 2018;25: 885–893. doi:10.1038/s41594-018-0124-7

16. Vilborg A, Passarelli MC, Yario TA, Tycowski KT, Steitz JA. Widespread Inducible Transcription Downstream of Human Genes. Mol Cell. 2015;59: 449–461. doi:10.1016/j.molcel.2015.06.016

17. Hennig T, Michalski M, Rutkowski AJ, Djakovic L, Whisnant AW, Friedl M-S, et al. HSV-1-induced disruption of transcription termination resembles a cellular stress response but selectively increases chromatin accessibility downstream of genes. Kalejta RF, editor. PLOS Pathog. 2018;14: e1006954. doi:10.1371/journal.ppat.1006954

18. Batra R, Stark TJ, Clark E, Belzile JP, Wheeler EC, Yee BA, et al. RNA-binding protein CPEB1 remodels host and viral RNA landscapes. Nat Struct Mol Biol. 2016;23: 1101–1110. doi:10.1038/nsmb.3310

19. Jia X, Yuan S, Wang Y, Fu Y, Ge Y, Ge Y, et al. The role of alternative polyadenylation in the antiviral innate immune response. Nat Commun. 2017;8. doi:10.1038/ncomms14605

20. Zheng D, Wang R, Ding Q, Wang T, Xie B, Wei L, et al. Cellular stress alters 3′UTR landscape through alternative polyadenylation and isoform-specific degradation. Nat Commun. 2018;9. doi:10.1038/s41467-018-04730-7

21. Shepard PJ, Choi E-A, Lu J, Flanagan LA, Hertel KJ, Shi Y. Complex and dynamic landscape of RNA polyadenylation revealed by PAS-Seq. RNA. 2011;17. doi:10.1261/rna.2581711

22. Zhu Y, Wang X, Forouzmand E, Jeong J, Qiao F, Sowd GA, et al. Molecular Mechanisms for CFIm-Mediated Regulation of mRNA Alternative Polyadenylation. Mol Cell. 2018;69: 62–74.e4. doi:10.1016/j.molcel.2017.11.031

23. Smith IL, Hardwicke MA, Sandri-Goldin RM. Evidence that the herpes simplex virus immediate early protein ICP27 acts post-transcriptionally during infection to regulate gene expression. Virology. 1992;186: 74–86. Available: http://www.ncbi.nlm.nih.gov/pubmed/1309283

24. Tang S, Patel A, Krause PR. Herpes simplex virus ICP27 regulates alternative pre-mRNA polyadenylation and splicing in a sequence-dependent manner. Proc Natl Acad Sci. 2016;113: 12256–12261. doi:10.1073/pnas.1609695113

25. Meier UT. RNA modification in Cajal bodies. RNA Biology. Taylor and Francis Inc.; 2017. pp. 693–700. doi:10.1080/15476286.2016.1249091

26. Pai AA, Baharian G, Pagé Sabourin A, Brinkworth JF, Nédélec Y, Foley JW, et al. Widespread Shortening of 3’ Untranslated Regions and Increased Exon Inclusion Are Evolutionarily Conserved Features of Innate Immune Responses to Infection. PLoS Genet. 2016;12. doi:10.1371/journal.pgen.1006338

27. Bentley DL. Rules of engagement: co-transcriptional recruitment of pre-mRNA processing factors. Curr Opin Cell Biol. 2005/05/20. 2005;17: 251–256. doi:S0955-0674(05)00048-7 [pii]10.1016/j.ceb.2005.04.006

28. Dubbury SJ, Boutz PL, Sharp PA. CDK12 regulates DNA repair genes by suppressing intronic polyadenylation. Nature. 2018;564: 141–145. doi:10.1038/s41586-018-0758-y

29. Krajewska M, Dries R, Grassetti A V., Dust S, Gao Y, Huang H, et al. CDK12 loss in cancer cells affects DNA damage response genes through premature cleavage and polyadenylation. Nat Commun. 2019;10. doi:10.1038/s41467-019-09703-y

30. Cortazar MA, Sheridan RM, Erickson B, Fong N, Glover-Cutter K, Brannan K, et al. Control of RNA Pol II Speed by PNUTS-PP1 and Spt5 Dephosphorylation Facilitates Termination by a “Sitting Duck Torpedo” Mechanism. Mol Cell. 2019;76: 896–908.e4. doi:10.1016/j.molcel.2019.09.031

31. Kecman T, Kuś K, Heo DH, Duckett K, Birot A, Liberatori S, et al. Elongation/Termination Factor Exchange Mediated by PP1 Phosphatase Orchestrates Transcription Termination. Cell Rep. 2018;25: 259–269.e5. doi:10.1016/j.celrep.2018.09.007

32. Huang K-L, Jee D, Stein CB, Elrod ND, Henriques T, Mascibroda LG, et al. Integrator Recruits Protein Phosphatase 2A to Prevent Pause Release and Facilitate Transcription Termination. Mol Cell. 2020;80. doi:10.1016/j.molcel.2020.08.016

33. Dai-Ju JQ, Li L, Johnson LA, Sandri-Goldin RM. ICP27 Interacts with the C-Terminal Domain of RNA Polymerase II and Facilitates Its Recruitment to Herpes Simplex Virus 1 Transcription Sites, Where It Undergoes Proteasomal Degradation during Infection. J Virol. 2006;80: 3567–3581. doi:10.1128/jvi.80.7.3567-3581.2006

34. Fraser KA, Rice SA. Herpes Simplex Virus Immediate-Early Protein ICP22 Triggers Loss of Serine 2-Phosphorylated RNA Polymerase II. J Virol. 2007;81: 5091–5101. doi:10.1128/jvi.00184-07

35. Galluzzi L, Yamazaki T, Kroemer G. Linking cellular stress responses to systemic homeostasis. Nature Reviews Molecular Cell Biology. Nature Publishing Group; 2018. pp. 731–745. doi:10.1038/s41580-018-0068-0

36. Sekulovich RE, Leary K, Sandri-Goldin RM. The herpes simplex virus type 1 alpha protein ICP27 can act as a trans-repressor or a trans-activator in combination with ICP4 and ICP0. J Virol. 1988;62: 4510–22. Available: http://www.ncbi.nlm.nih.gov/pubmed/2846867

37. Ramírez F, Dündar F, Diehl S, Grüning BA, Manke T. deepTools: a flexible platform for exploring deep-sequencing data. Nucleic Acids Res. 2014;42: W187–91. doi:10.1093/nar/gku365

38. Robinson MD, McCarthy DJ, Smyth GK. edgeR: A Bioconductor package for differential expression analysis of digital gene expression data. Bioinformatics. 2009;26: 139–140. doi:10.1093/bioinformatics/btp616

39. Quinlan AR, Hall IM. BEDTools: a flexible suite of utilities for comparing genomic features. Bioinformatics. 2010;26: 841–842. doi:10.1093/bioinformatics/btq033

40. Li H, Handsaker B, Wysoker A, Fennell T, Ruan J, Homer N, et al. The Sequence Alignment/Map format and SAMtools. Bioinformatics. 2009;25: 2078–2079. doi:10.1093/bioinformatics/btp352

